# iTome Volumetric Serial Sectioning Apparatus for TEM

**DOI:** 10.1101/2024.07.02.601671

**Authors:** David R. Peale, Harald Hess, P. R. Lee, Albert Cardona, Davi D. Bock, Casey Schneider-Mizell, Richard D. Fetter, Wei-Ping Lee, Camenzind G. Robinson, Nirmala Iyer, Claire Managan

## Abstract

An automated ultra-microtome capable of sectioning thousands of ultrathin sections onto standard TEM slot grids was developed and used to section: a complete *Drosophila melanogaster* first-instar larva, three sections per grid, into 4,866 34-nm-thick sections with a cutting and pickup success rate of 99.74%; 30 microns of mouse cortex measuring roughly 400 um x 2000 um at 40 nm per slice; and a full adult *Drosophila* brain and ventral nerve column into 9,300 sections with a pickup success rate of 99.95%. The apparatus uses optical interferometers to monitor a reference distance between the cutting knife and multiple sample blocks. Cut sections are picked up from the knife-boat water surface while they are still anchored to the cutting knife. Blocks without embedded tissue are used to displace tissue-containing sections away from the knife edge so that the tissue regions end up in the grid slot instead of on the grid rim.

## INTRODUCTION

Microstructural characterization of tissues has relied heavily on the high spatial resolution images produced via electron microscopy. Consequently, sample preparations have typically included the use of ultra-microtomes run by skilled artisans to produce the thin (<50nm) sections required for such imaging. Volume reconstruction of such microscopic detail is pushing the requirements on specimen preparation to ever larger numbers of sections and ever higher and more consistent quality of the sections to be imaged. Consequently, this is pushing the limits of human endurance while manually operating ultra-microtomes.

Within the last decade, a few new automated techniques have been developed to deal with this challenge, some employing microtomes, and some not. In one such approach, named BSEM, the entire sample block is mounted in an SEM, and the top-most layer is destructively removed by a microtome blade thereby leaving a fresh layer of block face to be imaged by the SEM (Denk and Horstmann, 2004). In another in-vacuum approach, named FIB-SEM, the face of the sample block is removed not by a mechanical blade, but by a focused ion beam, and the fresh sample face is again imaged by SEM (Knot et al. 2008; Xu et al. 2017). In another technique, named ATUM, a conventional ultra-microtome has been outfitted with a reel-to-reel tape pickup mechanism which synchronously drags floating sections out of the knife-boat water onto the tape as they are being cut. The tape is subsequently cut into strips which are mounted on silicon wafers, and the wafers are placed into a SEM for imaging (Shalek et al. 2011). A more recent development, GridTape TEM, etches slots into the tape and covers them with a strong support film, to then image floating serial sections with transmission EM (Graham et al. 2019).

In each of these methods except GridTape TEM, the block faces or sections are imaged in an SEM. While SEM imaging methods and speeds have been improving, TEM (specifically TEMCA; Bock et al. 2011; and its modern sibling, GridTape TEM; Graham et al. 2019) retains a significant speed advantage over SEM by virtue of the parallel nature of its imaging process. Such speed advantages greatly reduce the enormous time required to image a significant volume at nanometre resolution. Furthermore, tilting the grids in the TEM can provide additional structural information within the thickness of a single section via tomographic methods. Consequently, an automated method for producing large numbers of high-quality TEM sections would represent a significant step forward towards the goal of imaging and reconstructing significant volumes of anatomically functional tissue.

## RESULTS AND DISCUSSION

We present here a system for automated section cutting and pickup that is compatible with TEM requirements, and we demonstrate the viability and versatility of this approach by performing sectioning on several large samples. In a first example, a whole *Drosophila melanogaster* first instar larva was sectioned into 4,866 sections. The sections were 34nm thick, and placed 3 to a grid on 1,622 grids. The section pickup success rate during this process was 99.74%. In a second example, a volume of mouse cortex measuring 400 um x 2000um x 30 um thick was sectioned into 750 sections each 40 nm thick. In this case the sections were cut and picked up two per grid with the long axis of the sections parallel to the knife edge. In a third example, an entire adult *Drosophila* brain and connected ventral nerve column (VNC), a tissue volume measuring 400 um x 800 um x 344 um, was completely sectioned into 9,300 sections, each 37 nm thick. The pickup success rate during this process was 99.95%.

The cutting of each of these samples brings with it a particular significance. The cutting of the whole *Drosophila* larva represents the first time the entire larva body has been completely sectioned for TEM imaging, and will enable resolving the identity of all motor neurons and somatosensory neurons by tracing their axonal projects to and from the central nervous system.. The cutting of mouse cortex shows that significant large volumes of neural tissue are accessible to TEM imaging methods. The cutting of the complete adult *Drosophila* brain and connected VNC represents the first time that both the complete brain and connected VNC have been sectioned together.

During the cutting of the *Drosophila* larva, only 12.5 tissue sections were not placed within the grid slot area. This resulted in a successful sectioning and pickup performance rate of 99.74%. Of the 12.5 sections involved in the error events, 6.5 were errors directly attributable to human error as five cut sections were lost after failing to keep a water reservoir full, and 1.5 sections were lost due to grid film debris on a pickup grid. Unprovoked errors caused the loss of 6 sections: 3 were lost when one section train broke apart during pickup, and 3 sections were lost when one “pusher” section failed to cut.

During the cutting of the *Drosophila* brain and VNC, only 5 tissue sections were not placed within the grid slot area. This resulted in a successful sectioning and pickup performance rate of 99.95%. Of the 5 sections involved in the error events, 1 error early in the cutting run (section #144 which was not yet into tissue) was the result of human error when a typographical error was made during an adjustment to the section cutting speed. A second section (#802) was lost when a water handling pump broke down. That failure automatically paused the cutting run, and the pump was replaced. Cutting was successfully resumed about 5 hours after the initial failure of the pump. The third lost section (#1453) was lost because the base of the section failed to stay anchored to the knife edge at the end of the cutting stroke. The fourth section that failed to be picked up (#1815) resulted after the section cut thin immediately following a grid cassette swap. The fifth section lost (#4910) was lost after a servo actuator broke down and the cutting run was again automatically paused. Upon restarting the cutting roughly four hours later, the first section to cut was too thin near the knife edge and did not remain stably anchored to the knife for pickup.

The system, which we refer to as the iTome (short for interferometric microtome), differs in several significant ways from conventional ultra-microtome designs and operation which contribute to its ability to section for TEM. We outline these innovations briefly here before going into more detail below.

First, multiple sample blocks are held on a precision linear translation stage which provides a vertical slicing motion (Figure 1). This stage allows us to mount and cut more than one sample block at a time. When tissue dimensions are relatively small, it is desirable to cut several sections to a single grid. In this case, we mount a second block of resin without embedded tissue in it on the same slicing stage, and cut a section of “empty” resin (“empty block” or “pusher block”) after the desired N sections of tissue for one grid have been cut. The empty resin section, which we refer to as the “pusher” section, displaces the tissue sections away from the knife edge so that the tissue will be positioned in the filmed grid slot, and not on the metal rim of the grid (Figure 4).

**Figure 1.**
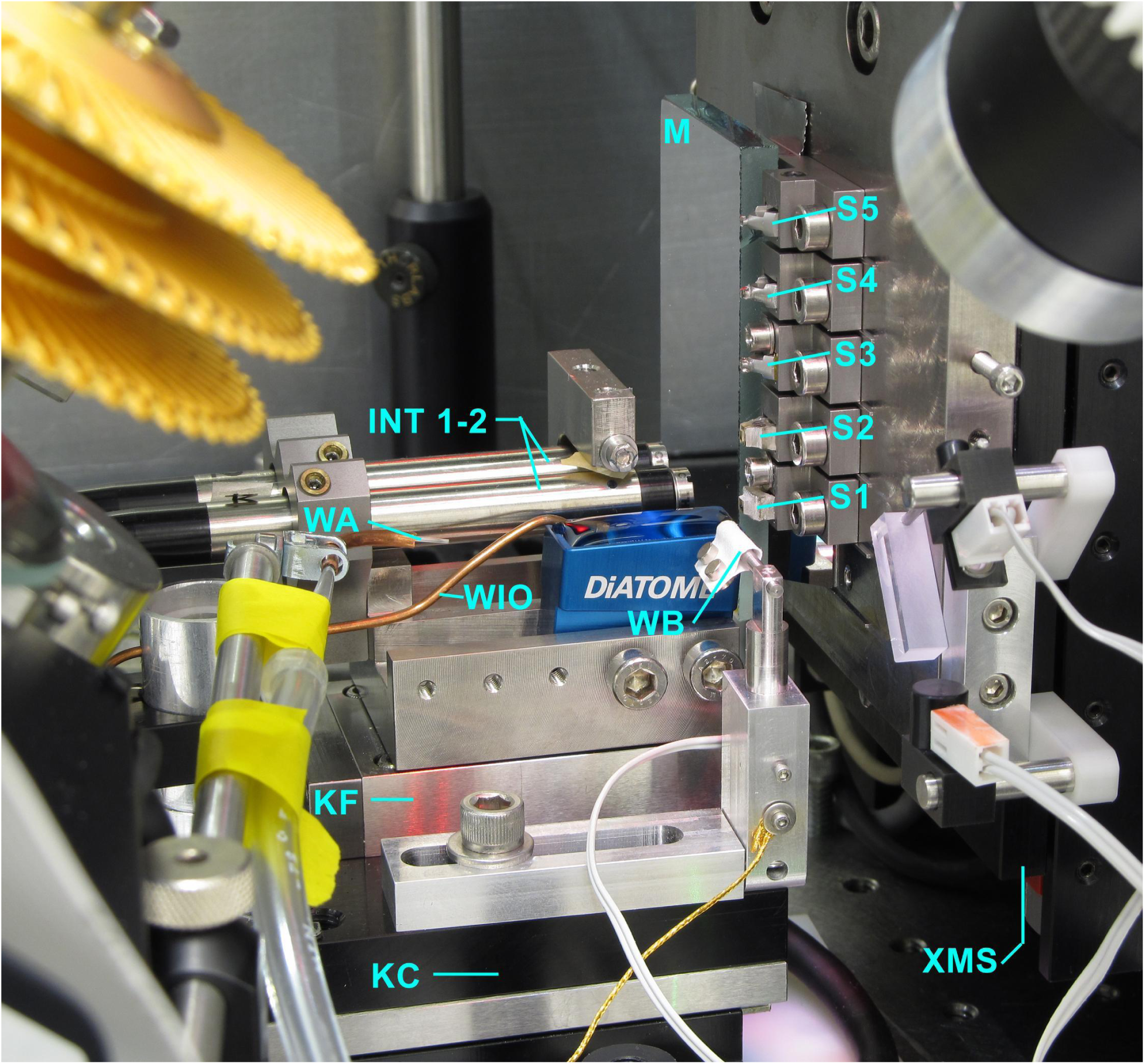
Principle components of the iTome involved with the knife and sample positioning and cutting operations. WB: knife-boat (also known as water boat); WIO: fill tube (water input/output); WA: vacuum aspirator tube (water aspiration); KF: stage with a piezoeletric linear actuator that controls the fine position of the knife; KC: a linear stage driven by a ball screw that controls the coarse-scale (micron to 25 mm) position of the knife; XMS: vertical linear motion stage (X-axis motion stage); S1 through S5: sample blocks (up to 5); M: mirror that serves as a reference surface which, in conjunction with fiber-fed interferometers “INT 1” and “INT 2”, allow us to measure and servo-regulate the distance between the sample face and the knife edge.

When tissue dimensions are large (e.g. 1000 x 2000 um) we cast the pusher section into the tissue block resin as one integrated block. In this case, we take one slice from the block per grid, but again, the pusher section ensures that the tissue will be positioned far enough from the knife edge to be in the open slot of the grid and not on the metal rim. As explained below, because we can accurately reproduce the knife-to-block distance, we are free to move the knife-boat after one slice as we execute the pickup motion, and return it to the proper cutting position for the next slice.

Second, optical interferometers measure a reference distance between the knife-boat and each sample block. This allows us to cut sections from either the tissue block or the empty pusher block as desired. This also reduces the sensitivity of the instrument to environmental perturbations and compensates for non-reproducible aspects of the mechanical positioners such as drive-screw backlash and piezo hysteresis. Since we can keep track of the cutting distance for each block face independently, we can precisely re-establish the cutting positions for either the tissue-containing or “pusher” blocks.

Third, section pickup is achieved while the section (or section train) is still anchored to the knife edge, not with the section(s) free-floating in the water as in manual ultramicrotome operation. Rather than touching and anchoring a corner or edge of the section to the grid face, and then dragging the section(s) out of the water as in conventional pickup methods, the iTome pushes the grid from under the floating end of the section in a forward and upward motion causing the free end of the section to slide freely up the wet grid face (see Suppl. Video). Water then drains from the grid and the section lies down onto the grid film without folds or wrinkles. With the sections now firmly attached to the grid film, the grid can then be further maneuvered to lift the pusher section off of the knife edge.

With these basic concepts outlined, we now present in further detail how they have been implemented.

The iTome consists of several key components and numerous other support features. Figure 1 illustrates some of the key components responsible for positioning the knife-boat and samples for cutting and section-pickup operations.

Multiple sample blocks “S1” through “S5” can be mounted on a vertical linear motion stage, “XMS”, which drives the slicing motion of the sample on the knife edge. Mirror “M” serves as a reference surface which, in conjunction with fiber-fed interferometers “INT 1” and “INT 2”, allow us to measure and servo-regulate the distance between the sample face and the knife edge. The interferometers use the reference mirror as a proxy for the actual sample face because it is impractical to use the actual sample face as an optical reference surface. The reference mirror is therefore mounted as close as practically manageable to the actual sample blocks in order to achieve a mechanically and environmentally robust connection to the samples. Likewise, the interferometers are mounted as close as manageable to the knife-boat to likewise assure a robust mechanical and environmental connection to the knife edge. Because the interferometers are offset to the side of the knife-boat, two interferometers are used in conjunction to compensate for any non-repeatable yaw in the motion of the stages that move the knife-boat towards and away from the samples. Because the interferometers were placed at the same height as the knife edge, there is no need to compensate for any non-repeatable pitching of the stages that move the boat. An acrylic enclosure also shields the apparatus from ambient temperature and humidity fluctuations which could affect the optical and mechanical paths linking the interferometers to the samples and reference mirror.

The knife-boat is a standard DiATOME Ultra 35-degree diamond knife. The knife-boat and the two interferometers are mounted on top of two positioning stages that control the advance of the knife and interferometers toward the sample blocks and reference mirror. Stage KF is a piezoelectric linear actuator that controls the fine-scale (nm to micron) position of the knife, and stage KC is a linear stage driven by a ball screw that controls the coarse-scale (micron to 25 mm) position of the knife. The coarse stage also serves to drive the motion of the knife-boat during section pickup and water cleaning.

Figure 2 shows the main components in the iTome that handle the TEM grids. A grid-holding cassette “GC” is made up of three stacked circular discs with 60 recessed pockets arranged circumferentially around each disc. The grids are retained in each pocket by a forked semi-circular cantilever spring that touches down on the outer lateral edges of the grid (See figure 3). The grid cassette is mounted on a rotary stage “GR” so that each pocket holding a grid can be turned to face manipulator tweezers “GT” that remove one grid from a pocket for section pickup. A separate linear stage “GL” lifts the cassette up and down along its axis to bring one of the three levels of the cassette into alignment with the height of the tweezers. When a desired grid pocket is facing the tweezers, and the tweezers have a grip on the grid, the cantilever spring for that grid can be lifted up by an actuator so that the grid can be freely removed from the pocket. Each grid thus picked is used for section pickup and then returned to the same cassette pocket for storage.

**Figure 2.**
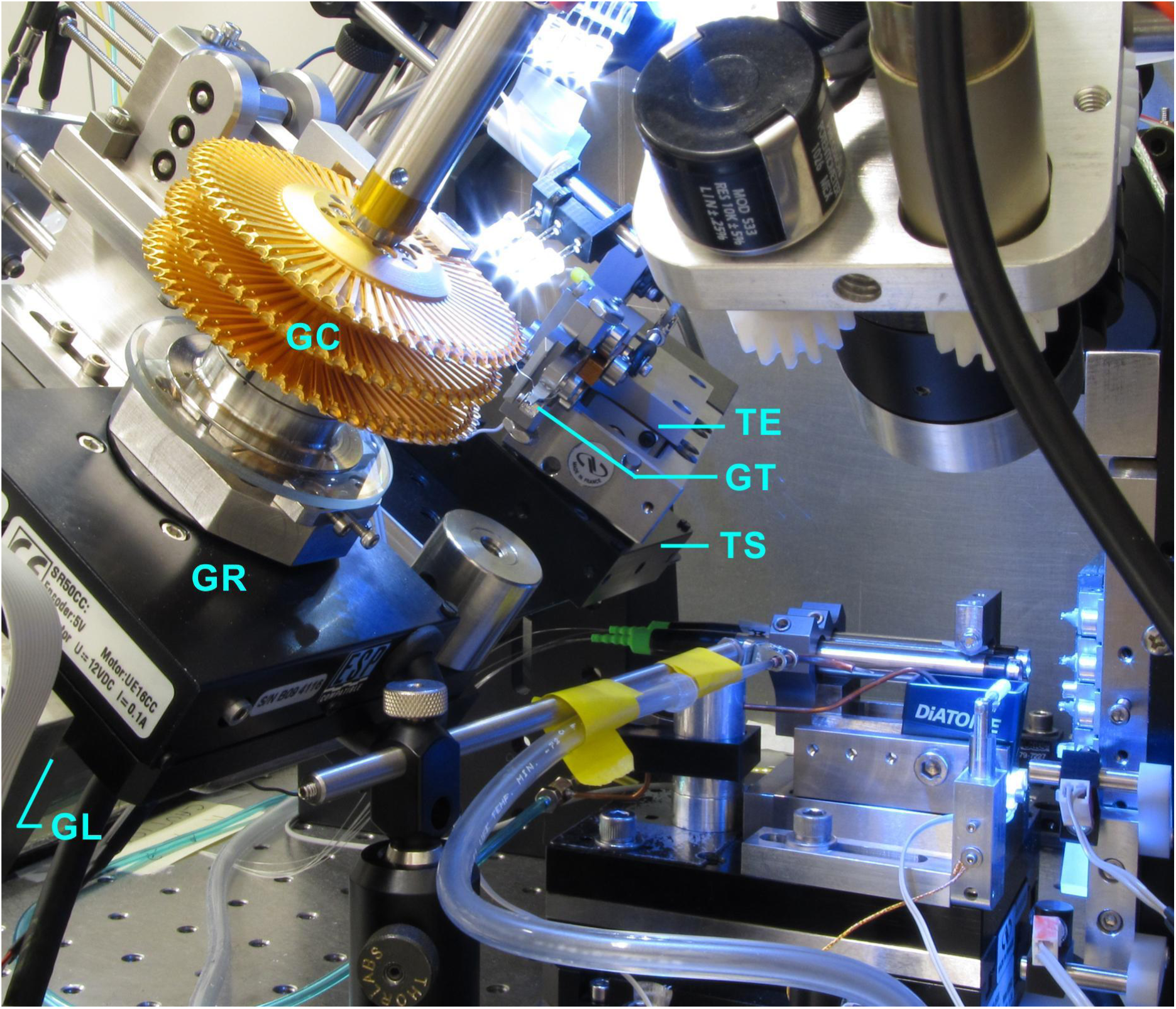
Principal components of the iTome involved with TEM grid handling. GC: grid-holding cassette, informally known as “daisies”; GR: rotary stage; GT: face manipulator tweezers to remove a single slot grid from the cassette for section pickup; TS: rotary stage of the GT manipulator tweezers; TE: linear-motion stage.

**Figure 3.**
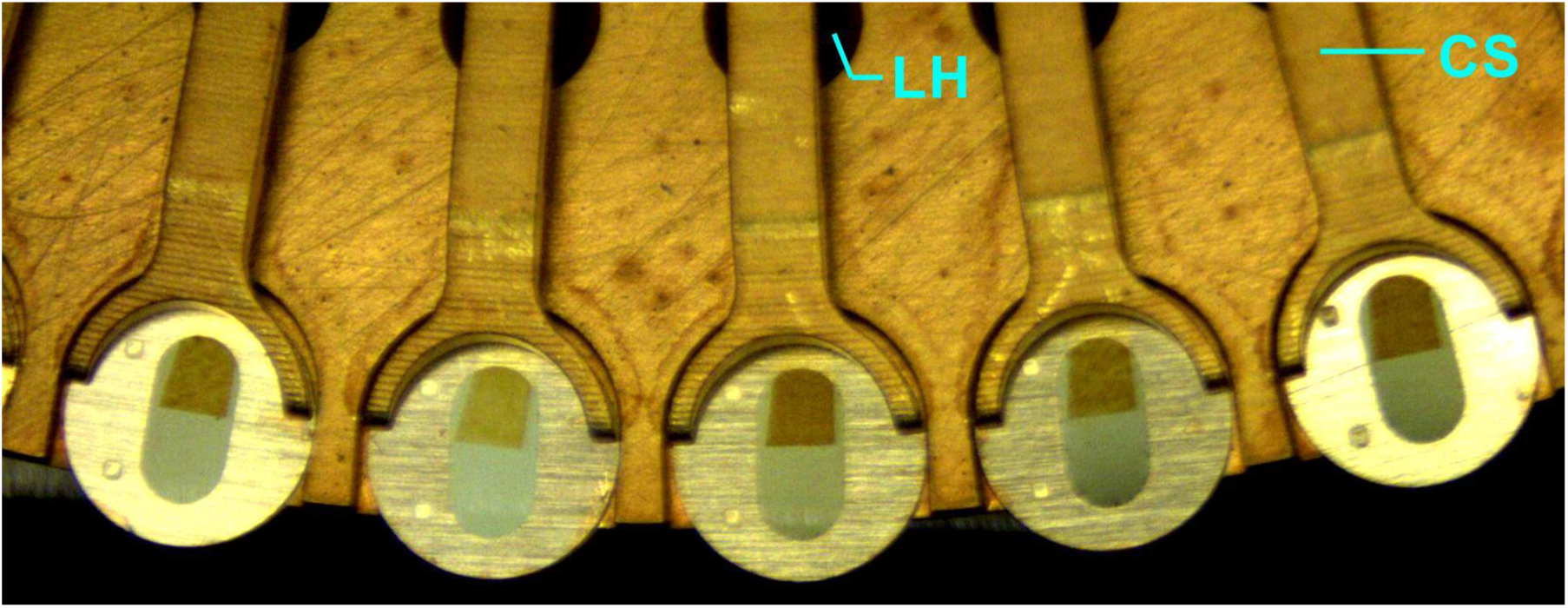
Close up view of the grid cassette pockets and grid retaining springs. A pin can be pushed up through hole “LH” to lift the retaining spring “CS” and allow a grid to be removed from the pocket.

**Figure 4.**
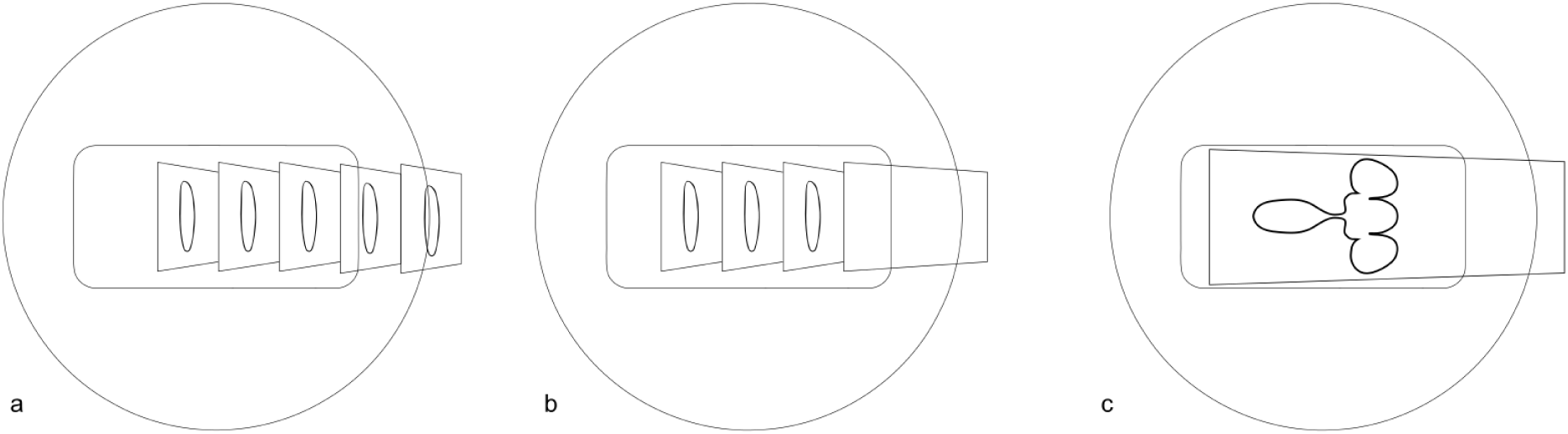
Outline at approximate scale of TEM slot grid and sections for iTome section pickup. 4a shows how two out of five tissue sections would overlap the metal frame of the slot grid if pickup were attempted with sections anchored to the knife without a “pusher” section. Because the iTome can address and cut multiple sample blocks in a random access fashion, multiple tissue sections can be cut, and a separate “pusher” section can be cut subsequently to move the valuable tissue sections away from the knife edge so that the grid can be maneuvered close enough to the knife edge to position the tissue sections in the open slot area. Figure 4b shows the arrangement of three tissue sections and one pusher section that can be produced in the iTome. Figure 4c shows a single section containing a large volume of tissue, and the length of “pusher” resin integral to the block that ensures that the tissue reaches the open slot area during section pickup.

The grid cassette is mounted via a dovetail base and can be removed easily and quickly from the iTome so that a replacement cassette with freshly prepared grids can be swapped into the iTome once the cassette in use has had all of its grids used for section pickup. This cassette swap, typically done once each day, takes about 60 seconds to perform, and therefore does not detrimentally disturb the machine’s equilibrium.

The manipulator tweezers “GT” are mounted on a rotary stage “TS” that allows the tweezers to swing from facing the grid cassette to facing in-line with the knife-boat. The rotary stage is inclined at an angle of about 37 degrees so that when the tweezers face the knife-boat, the grid is inclined to the water and can be extended forward and under the floating sections by a linear-motion stage “TE” that is mounted under the tweezers and on top of the rotary stage.

Figure 3 shows a close-up view of the edge of one level of the grid cassette. Once the grid-handling tweezers have a grip on a grid, the cantilever spring “CS” can be made to release its grip on a grid by pushing a pin up through hole “LH”. A video of the grid-handling and section-slicing operations can be seen in the supplementary materials.

Sections cut by the iTome are picked up from the water surface with 3mm-diameter TEM slot grids. These grids have roughly oval slots ∼ 2mm long by ∼1mm wide which are covered with pioloform films a few tens of nm thick.

In typical TEM section pickup, a skilled artisan will cut a number of sections from the block to form a train (or “ribbon”) of sections with the last section remaining anchored to the knife edge (Harris et al. 2006). A desired subset of these section(s) are displaced from the free end of the train using a hair or whisker and maneuvered, free-floating out onto the open water surface, away from the remaining sections. Holding the grid with tweezers in one hand and manipulating the free-floating sections with the hair in the other hand, the grid is inserted at an angle into the water, and maneuvered more or less under the sections. The grid is then lifted up, catching a corner of one section on the grid film and thereby dragging the section, or train of sections, up with the grid. The exact relative position of the grid with respect to the sections, the speed and smoothness of lift motion, and the hydrophilic character of the grid film are just a few of the crucial parameters contributing to the success and quality of the section pickup onto the grid film.

In contrast, the iTome cuts only the number of sections to be picked up on one grid at a time. In order to provide stability to the position of the cut sections during pickup, the last section cut remains anchored to the knife edge. If that section were a tissue section, then it would be impossible for the slot grid to be maneuvered in such a way as to place all of the tissue sections into the slot area of the grid. Figure 4a illustrates this predicament.

The iTome, by virtue of its ability to cut a section from any of the multiple blocks mounted on the slicing stage, solves this problem by cutting the last section from a second block of empty resin. This section is prepared to be long enough to displace the tissue sections far enough away from the knife edge so as to be in the slot area of the grid during section pickup. Figure 4b shows the arrangement of resulting sections on the grid produced by this method. In the case of large volume tissue samples, where one slice of tissue fills the majority of the 1×2mm grid slot, we cast and trim the tissue into a block such that there is a length of empty resin integral to the block to perform the “pusher” function. This arrangement is illustrated in figure 4c.

The section-pickup process is illustrated in Figure 5. We use a commercially available Ultra 35-degree diamond knife made by DiATOME. The profile of the knife-boat bottom and the angle of the diamond inside the boat constrain the available positions the grid can assume with respect to the section(s) while they are still anchored to the knife edge. However, instead of depending on the human artisan’s dexterity and rapid visual feedback to dynamically manipulate sections that have been detached and moved away from the knife edge, we perform section pickup while the last-cut section is still anchored to the knife edge and employ the “pusher” methodology described above.

**Figure 5.**
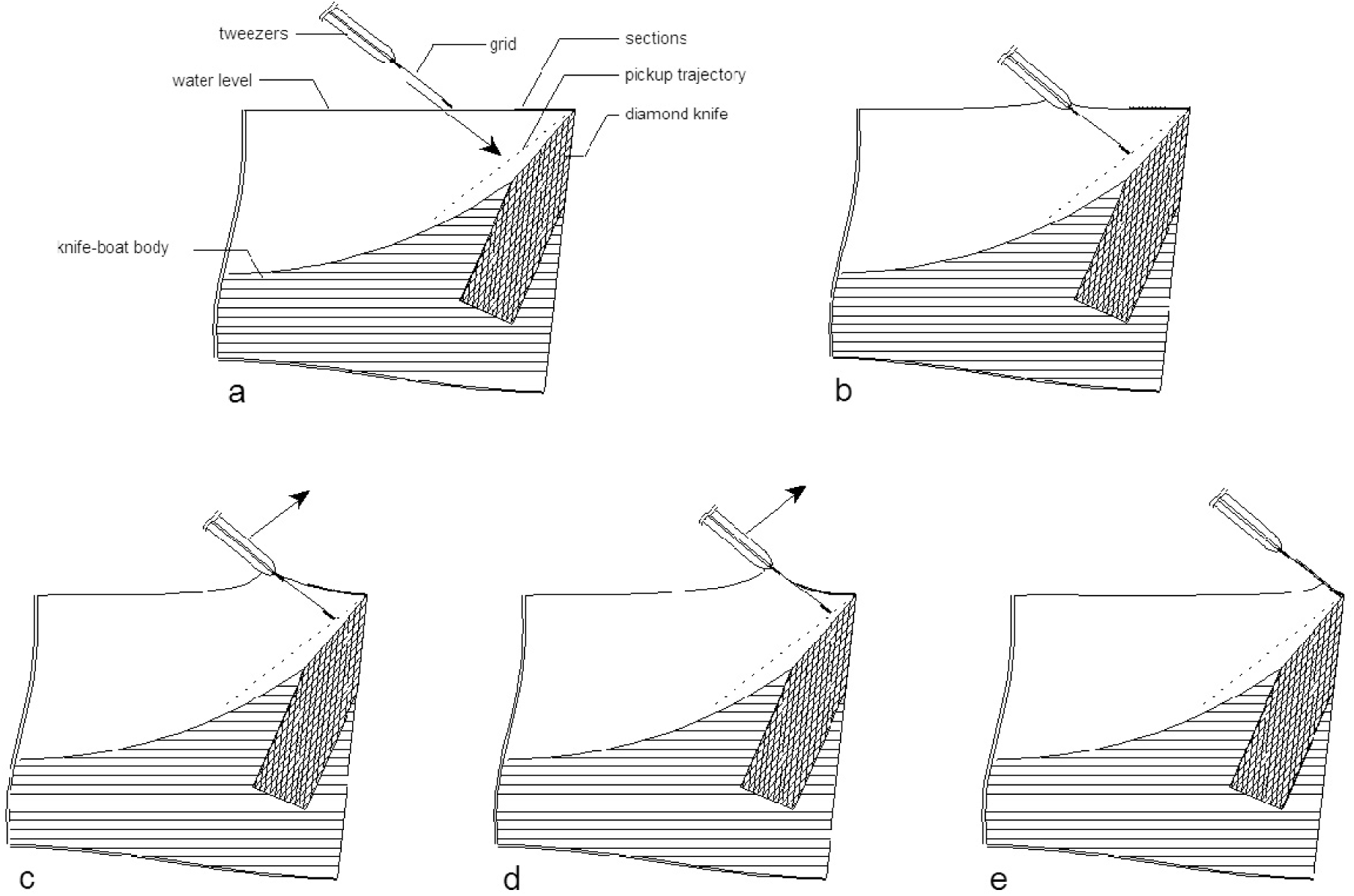
Approximate scale illustration of the region near the front of the knife-boat during the section pickup process. The profile of the boat bottom and angle of the diamond knife in the water constrain the trajectory available to the grid as it moves to pick up sections. Figures a and b illustrate the insertion of the grid into the water to the left of the free end of the section train to a depth such that the tip of the grid lies on the pickup trajectory line (shown as a dotted line below the water surface in the figures). Figures c, d, and e illustrate the motion of the grid during the section pickup operation. Between stages d and e, the curvature of the water meniscus on the face of the grid flattens. This causes the section train to slide up the face of the grid on the remaining water film. Once the water film drains completely, the section train is securely stuck to the grid film, and can be lifted off of the knife edge.

“Traditional” section pickup requires the grid to come up either directly from below the section or pulling up and back away from the section(s). These motions enable the grid film to contact some portion of a section and subsequently draw the section, or section train, up onto the grid film as it rises out of the water. Because of the constraints imposed by the boat-bottom geometry, these motions are not possible in the iTome. Consequently, the iTome takes advantage of the fact that the section, as long as it is floating on the water, can withstand compressive forces in the plane of the section that are much larger than gravity can impose. The gravitational forces on the section are significantly smaller than the forces arising from the difference in surface free energy between the water and the section. Because of this, it is possible for the grid to sweep up and forward into the section train and “push” the sections up the face of the grid while the knife edge supplies the necessary counterforce. Water then drains from the grid face, and the sections lie down flat and smooth and adhere to the grid film. Once the sections have been anchored to the grid film, the grid can be further maneuvered to lift the pusher section off of the knife edge.

One advantage of the iTome’s pickup process is that section placement within each grid slot is highly reproducible. Figures 6 and 7 each contain images of twelve consecutive grids from both the first instar *Drosophila melanogaster* larva, and the fly brain and VNC cutting runs. These images demonstrate the reproducibility of the section placement from grid to grid, both for multiple sections per grid and for one large section per grid. In the Instar-1 larva cutting run, each grid received 3 tissue sections which were pushed into the slot by one pusher section. In the Fly brain and VNC cutting run, each grid picked up one large section.

**Figure 6.**
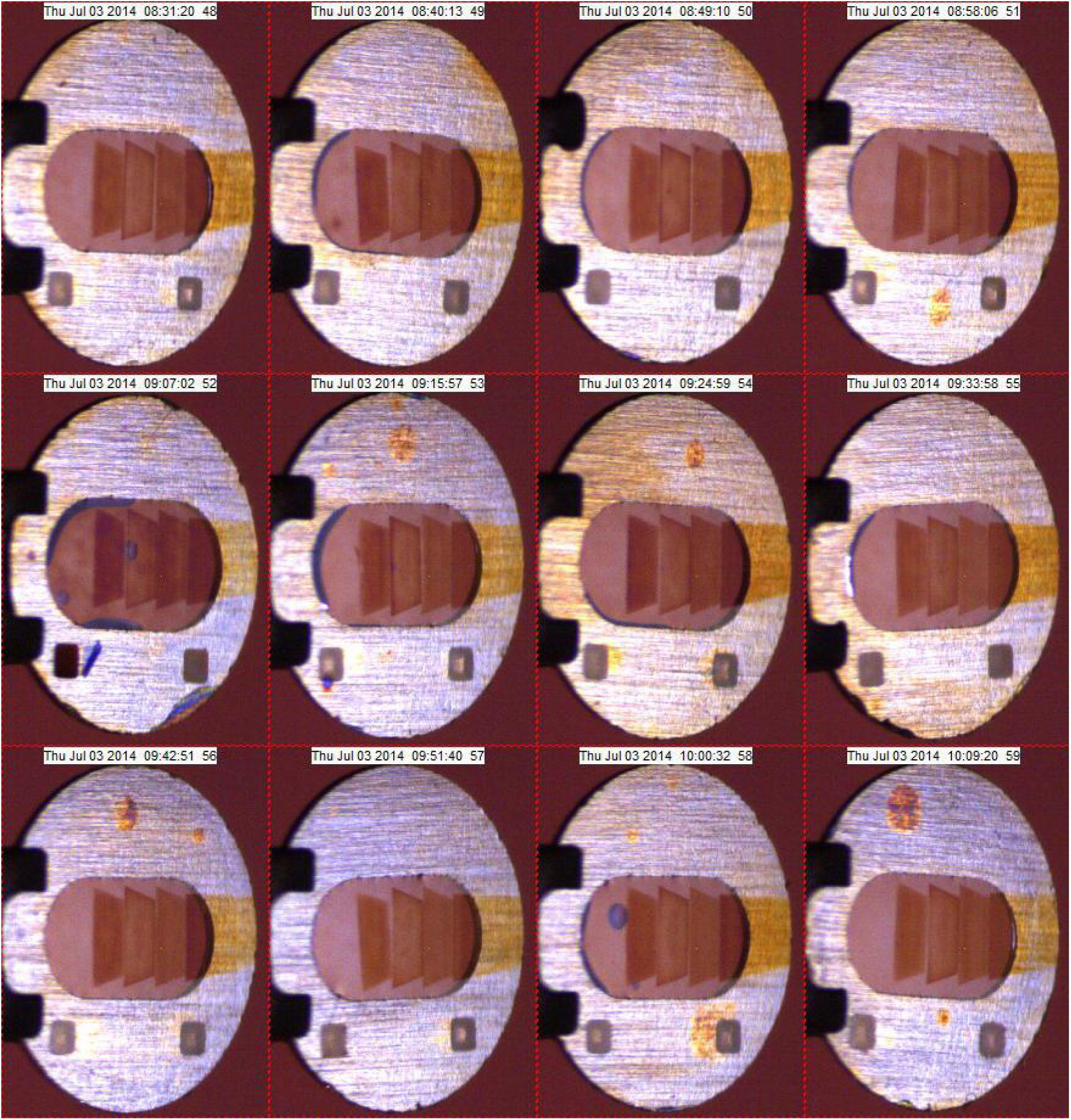
Images of 12 sequential grids from the Drosophila Instar-1 Larva cutting run shortly after section pickup, showing the grid-to-grid uniformity of the section placement achieved by the iTome section-pickup mechanism.

**Figure 7.**
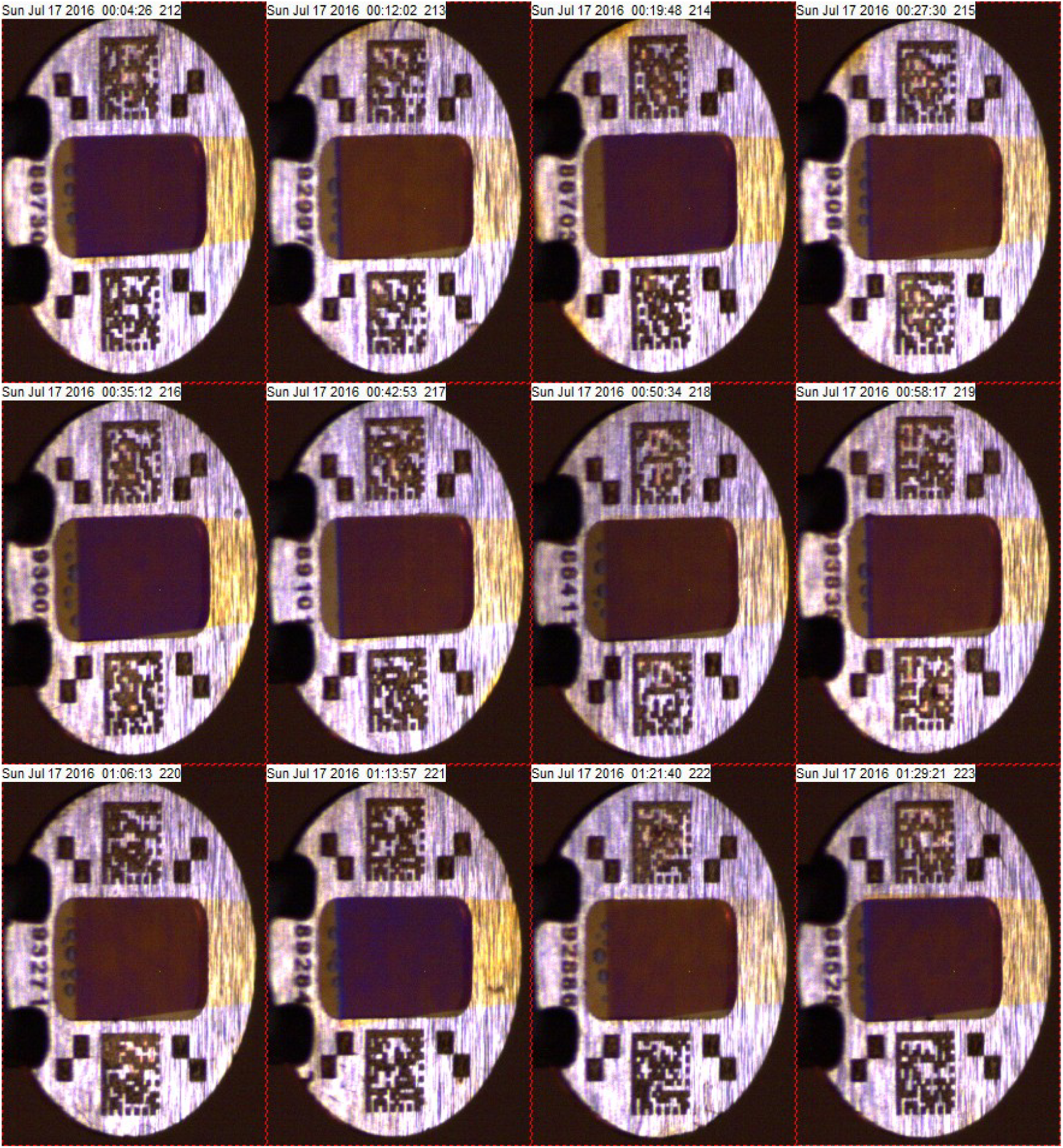
Images of 12 sequential grids from the fly brain and VNC cutting run shortly after section pickup, showing the grid-to-grid uniformity of the section placement achieved by the iTome section-pickup mechanism.

Such reproducibility is possible because we leave the section anchored to the knife during pickup. This enables us to laterally pre-position the grid to ensure that the section will be laterally well-centered in the slot after pickup. Furthermore, we also know ahead of time how long the pusher section needs to be in order to put the tissue sections in the longitudinal center of the slot. As long as we end the grid pickup motion with the grid edge at the same position w.r.t. the knife edge each time, the section will push up to the same position along the slot. Thus it is the simple static relation between the final relative positions between the grid and knife that determine where in the slot the tissue section is finally placed.

The iTome’s automated section pickup is in contrast to difficulties encountered in manual pickup, in which the multiple difficult-to-control parameters such as initial relative positions and angles of the section w.r.t. the grid, the exact speed and smoothness of the pickup motion, and the exact hydrophilic properties of the grid film play substantially into the final position of the section within the grid slot.

There are, however, dynamic factors that affect the section pickup quality and placement in the iTome. The grid film must remain wetted by the water during the pickup motion, or else the section cannot freely slide up the grid face. This puts a lower bound on both the speed of the grid pickup motion and the ability of the water to remain wetted to the grid film. Likewise, the viscous drag of the boat water on the section cannot exceed the in-plane compressive or shear strain stability limits of the section on the water, and/or the junctions between sections, and/or the anchoring strength of the section on the knife edge. This puts an upper bound on the speed of the pickup motion. Within these two limits however, highly reproducible pickup is achieved.

The dominant factors that affected the position of the sections within the grid-slots shown in Figure 6 were the angular orientation of the grids, and the angular yaw of the sections on the water at the time of pickup. The rotational orientation of the grids was determined when they were loaded into the grid cassette by a supporting robotic machine (separate from the iTome). We subsequently improved the angular accuracy of this automated loading step and reduced the slot angle fluctuations from grid to grid. The resulting improvement to the consistency of the grid slot orientation is visible in Figure 7. Secondly, some lateral yaw of the section train was occasionally produced during cutting as the last section was being parted from the sample block. This section-parting yaw is a function of many parameters relating to not only the block width, but detailed characteristics of the top surface of the block as well as the condition and uniformity of the knife edge doing the cutting. In manual pickup, large sections are more difficult to pick up without folds or wrinkles than small ones. In the iTome, the stability of the robotic mechanics and the nature and accuracy of the pickup method make picking up large sections no more difficult than small ones. Hence it has proven practical to mitigate the lateral section yaw by trimming the samples to be wide enough to provide a large baseline and thereby reduce the potential for lateral section yaw. We also pay close attention to the smoothness of the top surface of the sample block, and ensure that it is parallel to the knife edge so that one side of the section does not finish cutting before the other as this can lead to a transient yawing motion of the section train that can dislodge sections from each other and weaken the anchoring force of the pusher section to the knife. An improvement in the section yaw stability can be seen by comparing the sections in Figures 6 and 7.

The motions necessary to achieve the desired grid trajectory and final position w.r.t. the knife during pickup are supplied by the coordinated axial motion of the tweezers and the longitudinal motion of the boat. We use a Newport XPS motion controller and compatible Newport linear stages models MFA-CC and VP25-XA for the tweezers and boat motions respectively. The XPS controller is capable of slaving two stages together to execute such coordinated motions. In the example shown in Figure 5, a linear grid trajectory is shown, but a linear trajectory is not fundamentally required, and the XPS is capable of producing more complex slaved stage motions. We are free to employ the cm-scale motions of the knife-boat stage in the pickup operations because the interferometers track the distance between the boat-knife and the sample blocks. So we do not have to have a separate 2nd degree of freedom on the tweezers’ motion to achieve the needed pickup motion.

After sections have been picked up, excess water on the grid is removed by blotting the grid on filter paper that has been wrapped around a round rod which is supported next to the front of the knife-boat (indicated by label “WB” in Figure 1). By swinging the tweezers’ rotary stage a few degrees to the right of the section pickup-position, then extending the linear tweezers stage, and moving the knife-boat stage forward appropriately, the back side of the grid touches the filter paper, and the excess water clinging to the grid wicks into the filter paper. After a few tens of seconds of further air drying, the grid is dry and is returned to the grid cassette structure.

After a grid has been used for picking up sections, it is placed back into the pocket in the grid cassette from which it came. After the grids in the cassette have been used, the cassette is replaced with a cassette containing fresh grids. In the cutting runs described here, the used cassette was transferred to a supporting robotic machine which transferred each grid from each of the three levels of the cassette to a standard grid storage box that holds 100 grids (e.g. Ted Pella GSB100). From there, grids to be imaged were post stained with uranyl acetate and lead acetate.

For imaging of the Instar-1 larva sections, up to 60 stained grids were subsequently manually loaded back into a single-level version of the grid cassette assembly. This single-level assembly was then placed into a Zeiss Ultra 55 (modified for operation as a STEM) on a stage with six degrees of freedom so that any grid in the cassette can be placed into the beam for STEM imaging. The low profile design of the cassette disc allows it to take advantage of all 6 degrees of freedom in the beam. So the section could be tilted for tomography in addition to planar imaging if desired.

## SUPPORT FUNCTIONS

In order to keep the water in the knife-boat pristine during extended cutting operations, we use a small vacuum aspirator tube (“WA” in Figure 1) that is supported at a fixed position at the rear extreme of the knife-boat stage travel. In this way, the knife-boat stage can be backed up such that the surface of the water in the knife-boat touches the aspirator tube and water is drawn off. The aspirator tube is made of hydrophobic plastic and, as such, never effectively pierces the water surface, but instead effectively sucks only surface water into the tube.

The water for the knife-boat is provided through the fill tube (“WIO” in Figure 1) that is mounted so as to move forward and back with the knife-boat. This tube is a spur that tees into two tubes just behind the knife-boat, one of which provides fresh distilled water to the knife-boat, and the other of which drains to a waste-water flask. Water is moved via separate syringe pumps, one for filling and one for draining. The water level in the boat is sensed optically. A video camera records the distribution of light reflected from the water surface and feeds the images to the computer control system. Software analyzes the distribution of reflected light in the image and determines the level of the water in the boat. The syringe pumps can then be commanded to set a water level according to the task at hand.

In addition to monitoring the water level in the knife-boat, machine-vision feedback is used in a number of other operations within the iTome system. Slight variations in the grid position that arise from factors such as wobble and eccentricity of the grid cassette, required clearance space around each grid in the cassette grid pockets, and slight variations in the grip of the tweezers holding each grid make it impractical to handle the grids by pre-programmed dead reckoning alone. We therefore use video cameras and image processing software to make corrections to the nominal mechanical stage positions during operations such as picking a grid from, and returning a grid to, the cassette pockets, and establishing the precise relative position of the grid with respect to the knife-boat.

In addition to providing feedback for fine mechanical guidance, the video cameras also make it possible to monitor section quality, and create image logs of the operational process. An example of section-quality inspection is shown in figure 8 where images of two consecutive grids are shown. In figure 8a) and b), the grid-inspection illumination is set for brightfield (i.e. the white LED light is diffused, and the reflection from the grid surface is specular into the numerical aperture, N.A., of the camera lens). This illumination is suited for viewing the grid film and section thicknesses, and as well as the placement of the sections in the grid slot. In both of these images, the section pickup process appears to have gone well. In figure 8c) and d), the illumination has been set to darkfield (i.e. the red LED light is configured as a point source, and the specular reflection of this light is angled to fall just outside of the N.A. of the camera lens). In this case, light that has scattered off of discontinuities in the film is imaged. This reveals features such as holes in the film, particles on the film, the boundaries between the sections, and folds in the sections or film. Such a fold in a section can be seen in the left section of image 8d) and is marked by blue arrows.

**Figure 8.**
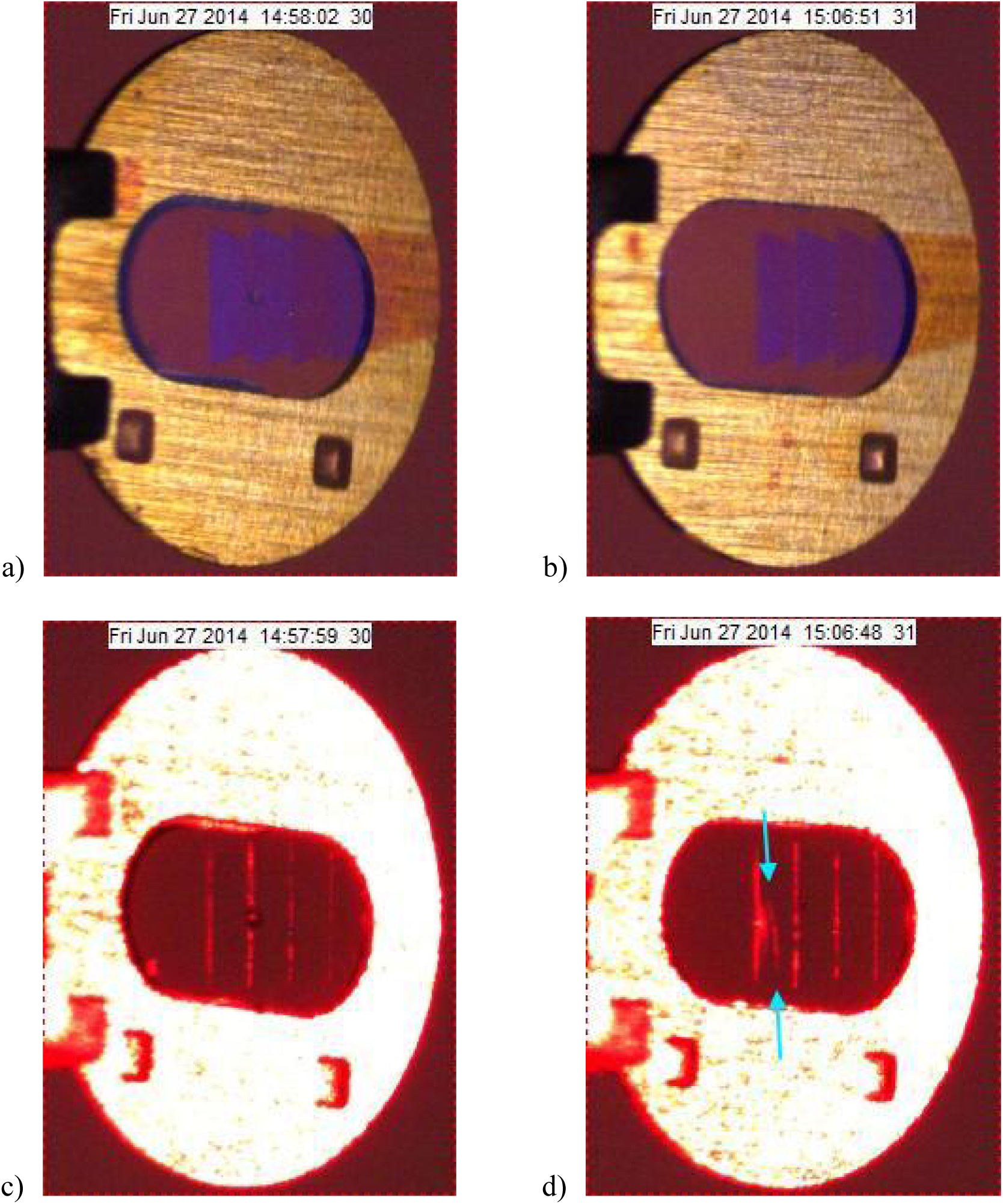
Images of two grids as reviewed in brightfield and darkfield illumination after section pickup. Images 8a and b show sections from two consecutive cutting cycles. In both cases, the section-pickup process appears to have gone well. Images 8c and d show the same two grids in darkfield illumination, which is sensitive to folds and other discontinuities in the section thickness. The boundaries between sections show up clearly, as does a fold in the left section of image 8d as marked by the blue arrows.

The ability to see such defects has provided useful feedback for adjusting the process conditions for optimal section pickup. In the case of the grid in figure 8d, other log images showed that the water had not fully wet the grid film high enough up the grid face, and the sections therefore “beached” prematurely onto the dry grid film. This resulted in the leading section “crumpling” slightly and producing folds. TEM imaging of that grid, shown in figure 9, shows the folds in exactly the location indicated by the darkfield image in figure 8d.

**Figure 9.**
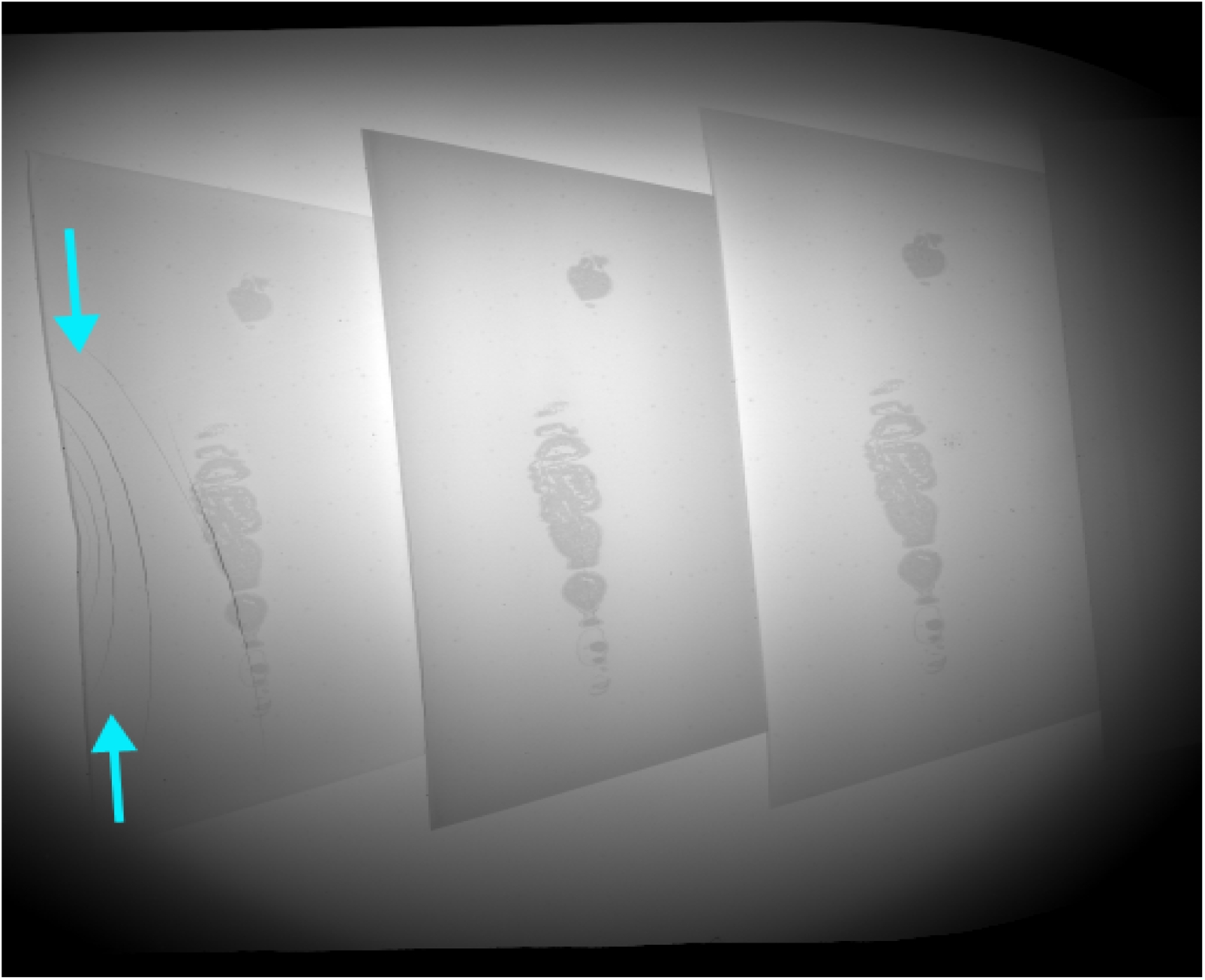
TEM overview image of the three sections on the grid shown in images 8b and 8d. The left section has several folds in it, as detected in the darkfield image 8d.

Thanks to such visual feedback, parameters for grid fabrication and section pickup can be adjusted to reduce the incidence of section folds. In the Instar-1 larva cutting run, a review of the TEM images at the scale shown in figure 9 showed that folds were present in the larva tissue in 52 (1.07%) of the 4866 sections, and more than half of those folds were present in the left-most section. The fold rate was subsequently reduced by simply inserting the grid deeper into the water before the section-pickup motion was executed.

Figure 10 shows STEM low-magnification overview images of a sequence of twelve contiguous grids, including the two grids shown in figures 7 and 8. The consistency and quality of the cutting and section pickup can be seen in these images. Again, the folds on grid 31 are marked by the blue arrows.

**Figure 10.**
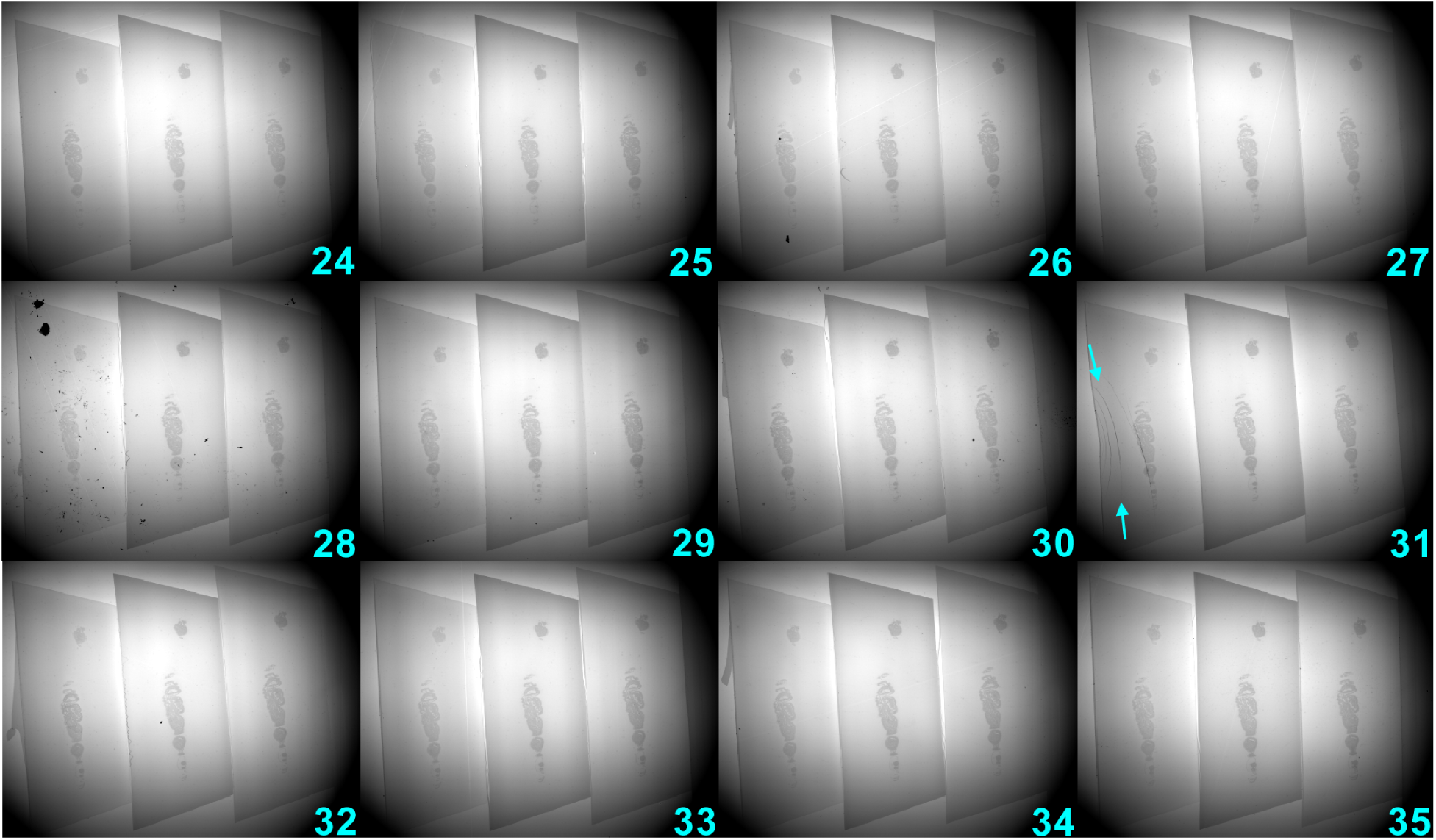
STEM overview images of 12 contiguous grids showing consistent cutting and pickup quality. Images labeled 30 and 31 are the same sections shown in figures 8 and 9.

**Figure 10.**
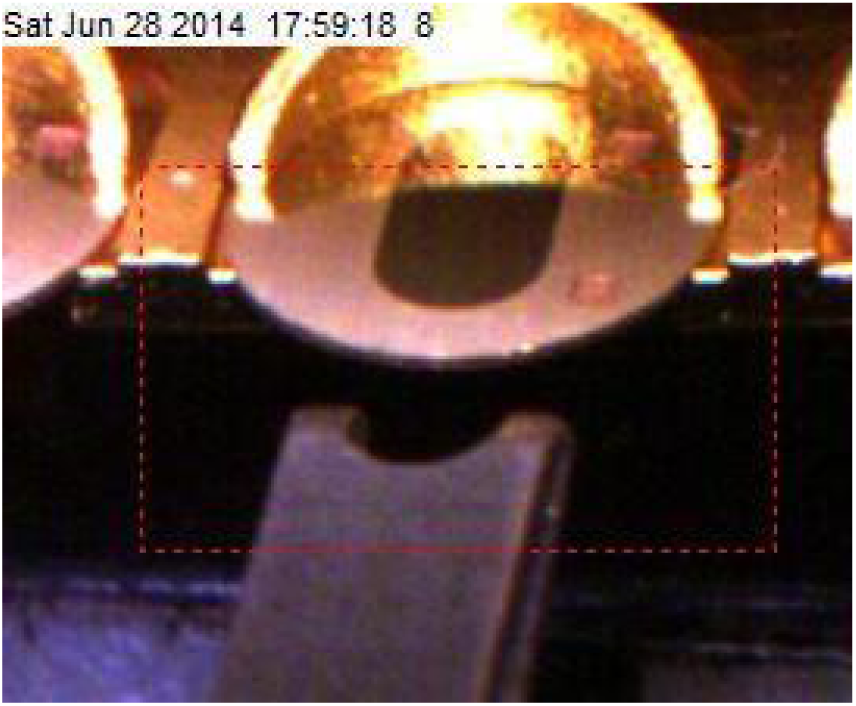
An image of the grid handling tweezers and a grid held by the cantilever spring in the grid cassette at the start of the grid-pick operation as produced by the machine vision camera that aids this process. The dotted red rectangle is the sub-image that is passed to the image processor to determine the exact relative position of the tweezers with respect to the grid slot.

The cameras also record other useful information during each cutting cycle. Before section pickup, the cameras make it possible to re-inspect the grid film for flatness, thickness (as indicated by color) and particulate contamination. We record images of the grid during insertion into the water, to monitor the proper wetting of the grid film, and of the water draining from the film, after the sections have been pushed up the wet grid surface. Finally, we record images from the cameras that monitor the picking and stowing of grids from, and back into, the grid cassette. This level of image logging facilitates tracking grids throughout the operation of the iTome and helps debug rare errors and further improve the operational reliability.

## Conclusion

We have designed and demonstrated a new process and apparatus for autonomously cutting and picking up ultra-thin sections on TEM slot grids. The cutting apparatus employs optical interferometers to regulate the knife-to-sample distance. This feature allows the apparatus to cut sections from more than one sample block in a random-access manner. The pickup process reliably places the cut sections in the open slot area of the TEM grid with a low incidence of folds or wrinkles. The pickup process exploits the ability of floating sections to be pushed up an inclined wet grid face while the sections are still anchored to the knife edge. We demonstrated the reliability of this system by continuously sectioning a whole Drosophila instar-1 larva into 4866 sections, each approximately 34 nm thick, and placing these sections, three to a grid, onto 1622 TEM grids over a period of less than 11 days.

This process and apparatus advance the goal of achieving high-resolution reconstructions of functional volumes of biological tissues by enabling the use of transmission electron microscopy for the imaging of the cut sections because the parallel nature of TEM imaging provides a throughput advantage over the scanned imaging methods required by the previously demonstrated autonomous sectioning methods. Furthermore, the ability to tilt the sections in the beam opens up the option of using tomographic methods for extracting further structural information from within the thickness of one section.

## Methods and Materials

### Stages and Functions

A Newport XMS100 linear stage carries the sample blocks. This stage has a linear electromagnetic motor and a glass scale for position feedback and is capable of 1-nm incremental motions over a range of 100mm. This stage is not intended for use in the vertical position, so a counterweight on a rocker arm is used to balance the static load of the vertical stage platform and the hardware mounted on it. In this configuration, the stage motor has to supply only enough force to accelerate the stage for motion, and a slight holding force when the rocker arm is tilted away from horizontal. This counterbalance setup minimizes the heating of the stage due to holding currents in the motor.

The knife-boat is mounted on top of a linear piezo stage (Physik Instrumente model PI-752.11C), which provides the fine motion used for servo control of the knife position during cutting. This piezo stage has an extension range of 35 um and minimum motion of 0.1nm. A PI E610 amplifier drives the piezo stage. Custom electronics drive the amplifier and interface it to the computer control system.

The piezo stage is mounted on top of a Newport VP-25XA linear stage which provides the coarse linear motion needed for operations such as section pickup, grid drying, and water cleaning. This stage is driven by a fine-pitch ball screw and has a travel range of 25 mm and an incremental motion of 100 nm. This stage was chosen in part because the ball screw provides a rigid drive mechanism so that when the stage is motionless during cutting, it does not need active feedback from the controller. Such feedback would have had a response time slower than the piezo stage in the servo feedback loop, and consequently, would have interfered with the stability of the servo loop.

A Newport PR50CC rotary stage provides the swinging motion of the tweezers between the grid cassette and the knife-boat. This stage has an incremental motion of 0.015 degrees and swings at a maximum of 20 degrees per second. A Newport MFA-CC linear stage is mounted on top of the rotary stage, and provides the extension motion needed for operations such as picking a grid from the cassette, dipping the grid into the boat water during section pickup, and stowing the grid back into the grid cassette. This stage has a travel range of 25 mm and an incremental motion of 100 nm.

A Newport SR50CC rotary stage provides the rotary motion to turn the grid cassette and align a grid pocket with the grid tweezers. This rotary stage has an incremental motion of 0.004 degrees, and a maximum rotation speed of 4 degrees per second. A Newport VP25-XA linear stage supports the rotary stage vertically so that the grid cassette can be moved up and down along its rotation axis to align any of the three cassette levels with the grid tweezers.

All of the above mentioned stages are interfaced to a Newport XPS motion controller. The XPS controller is capable of slaving two stages together to execute coordinated motions such as were needed for the grid pickup operations. In the example shown in figure 3, a linear grid trajectory is shown, and such a trajectory was used for the cutting of the full larva reported here, but a linear trajectory is not fundamentally required, and the XPS is capable of producing more complex slaved stage motions. A custom circuit that tied into auxiliary I/O lines available on the XPS controller detects electrical continuity between the grid-tweezers and any grounded object, including the grids in the grid cassette and the knife-boat water surface. The Newport XPS controller communicates with the main control computer (a quad core PC running Windows 7) via a direct Ethernet TCPIP connection. This provides a fast, low latency and reliable connection between the XPS and main control PC.

### Other Computer-Interfaced Hardware

In addition to the XPS, the control computer also communicates with several other peripherals. As mentioned previously, four video cameras (3 megapixel iDS model UI-1460SE-C) each use a USB 2.0 connection to feed video to the control software. All four cameras feed video at all times, each one to a separate standalone application that handles that video feed. Numerous LED sources supply light for the video cameras. These LEDs are driven by a custom sixteen-channel current supply and are interfaced to the main computer via a National Instruments PCIe-7854R card mentioned below. This LED control unit also provided up to eight pulse-width modulated (PWM) signals, four of which controlled rotary servo actuators (Model Airtronics 94761z). Three of these servos control the lifter arms that release the grid-pocket clamping-spring for each of the three levels of the grid cassette. The fourth servo controls the opening and closing of the grip of the tweezers that handled the grids. These servos have the favorable characteristic that the servo motor current is cut off when the PWM signal is set to 0% duty cycle. This prevents unwanted mechanical motion around the holding position, and the associated vibration created by such jitter. It also ensures a constant and nearly zero power dissipation. This feature contributes to extending the lifetime of the servos.

A serial port link communicates with two New Era NE-1000 syringe pumps that fill and drain the knife-boat water. Auxiliary outputs available on the NE-1000 pumps control a custom rotary valve that refills the source water syringe and empties the drain water syringe at the end of their respective strokes.

The optical interferometers were custom built by Integral Physics to Engineering, LLC, and interfaced to the computer via a National instruments field programmable gate array (FPGA) interface card, model PCIe-7854R. This card has eight simultaneous sampling 16-bit analog inputs, eight 16-bit analog outputs, and 96 digital I/O lines, all tied into a Virtex-5 LX50 FPGA which can be configured via Labview software to perform complex real-time acquisitions, computations and outputs. A direct memory access (DMA) interface to the host PC enables rapid data transfer from the card to the PC memory with minimal software intervention and overhead. This FPGA interface implements the servo control loop containing the interferometer sensing and the piezo actuator. The DMA link reports sample block position data (as sensed by the interferometers) and piezo output voltages (as commanded by the FPGA) to the host PC so that the main control program can monitor, graph and log the sample and piezo actuator positions.

### Control Software

Computer-control software of the entire iTome apparatus was written in Labview. Various aspects of the operations were broken into separate applications. The main application handled the interface to the mechanical stages, syringe pumps, servo actuators and LED illumination controls. The main application also handled the input, graphing and logging of interferometer and piezo-stage data from the FPGA card, and provided the configuration interface for setting the servo loop control parameters. It also contained a custom-written script interpreter so that text-based scripts could be written to direct operations of the various elemental functions of the system. This made it easy to encapsulate operations of the system into complex higher-level functions, and through these, to make changes in the operations of complex functions on a case by case basis. The main program also logged virtually every script command and user-interface state change via a UDP stream. A separate logging application received these data and stored them to disk. The logging application also filtered the command stream in real time to create a second log containing user-specified commands that were useful in monitoring cutting cycle operations (e.g. the date, time, grid number, cassette number, cassette level, and cassette pocket from which a grid was picked from or into which it was stowed, during a cutting cycle).

Each of the four video cameras is controlled by its own separate instance of a camera control and capture application. The main iTome program communicates with these video applications via a Labview “Shared Variable” protocol. Through that channel, script commands from the iTome main program set camera parameters such as exposure time, frame rate, CCD gain, pixel binning, and region of interest size. Similar commands trigger the capture of a frame of video for image analysis or simply cause an image to be saved to disk as part of operational log documentation.

Likewise, a script command from the main iTome program causes images from any of the four camera control applications to be passed to an image processor application. Another script command causes the image processor to perform a task-specific analysis. The image processor uses a variety of image-processing methods including thresholding, differential comparison, and correlative methods to determine the parameters of interest for each task-specific case. The resulting parameters from the image processor are passed back to the main iTome program for use in the scripting interpreter which can incorporate those results in flow-control branching decisions, or as parameters to adjust stage positions.

A separate application controls the water level in the knife-boat, and communicates with the main iTome program via Labview shared variables. The water-control program runs continuously, and periodically adds a quantity of water to the knife-boat to make up for evaporation. The program also responds to commands from the iTome program to pause the automatic-refill functions, or raise or lower the water level.

### Machine Vision Operations

As mentioned in the body text, the video cameras and image processing software are used in the iTome to assist in the picking and stowing of grids from and to the grid cassette, and the inspection of the grids, the grid slot orientation, grid film quality, and section placement in the grid slots during the course of a cutting cycle. Figure 10 shows an example image from one camera used to guide the picking and stowing of grids from and to the grid cassette. In this case machine vision makes corrections to the position of the tweezers because the cassette inevitably has a small amount of eccentricity and wobble to it that causes the position of the grids to wander as the cassette rotates through 360 degrees.

Machine vision is also used to establish a precise spatial relationship between the grid edge and slot and the knife edge once the grid has been picked from the grid cassette by the tweezers. This is necessary because it is the relative position of the grid with respect to the knife edge that determines the centering of the sections in the grid slot at the end of the pickup motion. The accuracy of the section placement in the slot is only as good as the accuracy of the grid-to-knife spatial relationship.

### Grid Loading Support Robot

When loading grids into a cassette, freshly prepared grids were loaded each day into a grid cassette by means of a robotic grid handling machine. The layout and hardware for this machine was similar to the iTome except that the slicing stage hardware was occupied by either a tray capable of holding 64 freshly prepared grids, or a grid storage box, depending on whether grids were being load into the cassette or being removed from it after cutting. Also, hardware to perform a grid rotation function unique to the grid handling machine was also present.

The hardware for the grid rotation operation consisted of a down-looking camera observing a 2 mm diameter rotary post. The purpose of this hardware was to allow the orientation of the grid slot to be set to a desired angle. Typically the desired slot orientation was with the major axis aligned along the grid cassette radial direction as can be seen in figure 3. For loading grids into the grid cassette, the grid tweezers would pick a grid from the fresh grid tray by dead reckoning and swing the grid to the rotary post where it would be put down on the post with the aid of machine vision from the down-looking camera. The same camera would then view the grid and the image processor would determine the slot orientation. The rotary post would then turn the grid slot to the desired orientation, and the grid tweezers would lift the grid up off the post and swing to the grid cassette where the grid would be placed in a pocket.

When it was time to unload grids from a grid cassette, after the grids have been used for section pickup and the cassette has been removed from the iTome, the grid supply tray was replaced by a standard grid storage box, e.g. Ted Pella style GSB100 or equivalent. In the unload operation, a grid was picked from the grid cassette, and the rotary post was optionally used to change the tweezers grip on the grid from aligned parallel to the slot to being aligned perpendicular to the grid slot. The grid was then lifted from the post and the tweezers swung to the grid storage box where the grid was inserted into the diamond shaped pocket of the box with the aid of a camera and machine vision. Figure 11 shows images of the grid about to be put into the box pocket, and the image processor feature detection results which guide a fine correction for the positions needed to insert the grid into the pocket without touching the pocket walls. Putting the grid into the storage box with the slot sideways lowers the risk of damaging the grid film with tweezers when the grids are hand-picked from the box later during staining.

**Figure 11.**
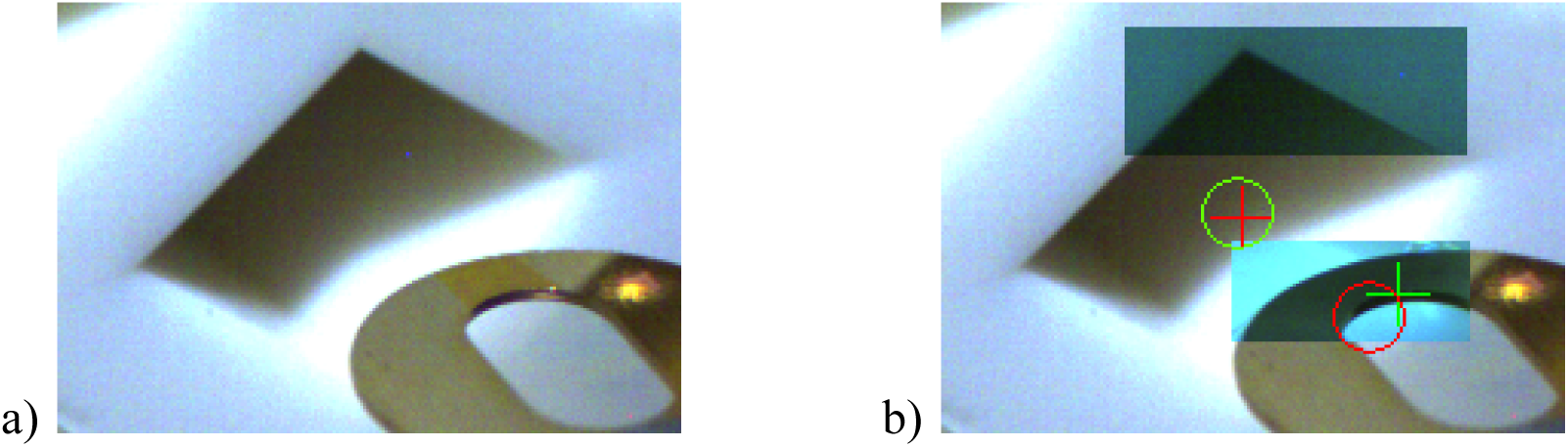
Images from the camera that guides the placement of grids into the storage box pocket in the grid handling robot that supports the iTome. Image a) is the input image to the machine vision image processor. Image b) shows the features of the pocket and grid which are matched by the image processor to make fine corrections to the tweezers position so that the grid is inserted into the pocket without touching the walls of the pocket.

## Acknowledgements

Dr. David Peale thanks Dr’s. Gleb Shtengle and Shan Xu for numerous instances of assistance with computer hardware and software. We thank HHMI Janelia Research Campus for funding.

## Supplemental Material

A movie showing many of the operational steps in the iTome during a cutting and section-pickup cycle. Please note that because of limitations in the video recording abilities from the cameras in the iTome, the operations shown are not all from one particular cutting cycle.

